# ALS mutations in the TIA1 RNA granule protein have differing effect on low complexity domain fibril formation

**DOI:** 10.64898/2026.05.18.725988

**Authors:** Yuuki Wittmer, Dylan T. Murray

## Abstract

Mutations in the low complexity domains of RNA-binding proteins are associated with neurodegenerative disease pathology. The TIA1 RNA-binding protein harbors seven such mutations linked to a clinical cohort of ALS patients. The altered low complexity domain sequence increases the number of TIA1-rich stress granules in cultured cells, delays their disassembly, and is associated with increased fibril formation. Altered molecular motions and contacts in condensed states like stress granules that result in the formation of amyloid-like fibril states is commonly observed for RNA-binding biomolecular condensates. Here we focus on the influence of the ALS mutations on fibril formation of the TIA1 low complexity domain. Repetitive seeding preparations of the seven TIA1 protein mutants all yield amyloid-like fibrils based on transmission electron microscopy images and increased thioflavin T fluorescence. Analysis of solid state nuclear magnetic resonance spectra recorded on all seven mutant fibrils reveals distinct structural differences in the relative to wild-type fibrils. Our results shed light on how the mutations affect structural conformations accessible to the TIA1 low complexity domain.

## Introduction

Heterogeneous nuclear ribonucleoproteins (hnRNPs) have been extensively studied for their wide variety of biological functions including mRNA splicing and translation. hnRNPs are linked to neurodegenerative pathology in amyotrophic lateral sclerosis (ALS), frontal temporal dementia (FTD), and Alzheimer’s disease.^1^ Many hnRNPs are RNA-binding proteins that contain a low complexity (LC) domain with a primary sequence compositionally biased to a subset of the 20 naturally occurring amino acids.^2^ Neurodegenerative disease associated mutations are concentrated LC domains.^1^ However, the conformational landscape for LC domains in relation to either diseased or functional states remains poorly characterized at high resolution. LC domains tend to be primarily disordered in solution, making functional and pathological conformations very challenging to study.^3^ Although approximately 30% of human proteins contain a low complexity domain, very few have been structurally characterized.^4^

Solid-state nuclear magnetic resonance (ssNMR) is a versatile tool for structural characterization, allowing for high resolution structure determination, characterization of protein motions, and providing a spectroscopic fingerprint useful for quickly tracking structural features and transitions^5^. A high-resolution structure for alpha synuclein fibrils related to Parkinson’s disease^6,7^ has been determined, featuring a Greek key topography with a hydrophobic core.^8^ Other structures include beta-amyloid fibrils related to Alzheimer’s Disease, also exhibiting a hydrophobic core^9,10^. More recently a structure was determined for fibrils formed by the LC domain of the RNA-binding protein fused in sarcoma (FUS) featuring the absence of a hydrophobic core and a less stable structure^11^. Although FUS is implicated in neurodegenerative disease, functional assays posit limited fibril-like association of the LC domain might be functional^11^.

Another hnRNP RNA-binding protein similar to FUS is the LC of the TAR DNA-binding protein 43 (TDP-43). TDP-43 LC forms amyloid-like fibrils in brain tissue of patients with ALS and FTD.^12,13^ Different morphologies of the TDP-43 LC fibrils have been characterized using both cryo-EM^13–15^ and solid state NMR^16,17^, revealing the presence of competing fibril cores with differing hydrophobic residue content contained withing a single LC domain. The FUS LC domain also contains two fibril-prone regions^11,18^. Although structural polymorphism in amyloid fibrils may be a general pathogenic feature, studies of LC domain fibrils suggest competing functional and pathogenic structural conformations that may partly be due to differing hydrophobic residue content.

The TIA1 RNA-binding protein contains a LC domain capable of forming fibrils and has a gradient of hydrophobic residue content. The TIA1-LC is involved in stress granule assembly and formation^12,19^, and contains seven ALS-linked point mutations.^1,12^ These mutations delay stress granule formation in cells, and alter in vitro liquid droplet biomolecular condensation by decreasing droplet phase fluidity and increasing partitioning into the droplet phase^20^. Importantly, these mutations tend to increase the propensity for TIA1 to form amyloid-like protein fibrils^20^. Droplet rigidification has been shown to be structurally distinct from fibril formation^21^. Thus, the TIA1-LC domain is a prime candidate to investigate the potential for another LC domain to balance pathogenic interactions driven by hydrophobic residues with polar functional interactions like TDP-43. The ALS-linked mutations are both near and distant to the fibril-prone region of the LC domain, raising the possibility that disease mutations tilt the balance of interaction in the TIA1-LC domain from functional to pathogenic. However, the TIA1 mutant fibril structures have not been structurally characterized.

Here, we report initial structural characterization for fibrils formed by each of the seven ALS-linked TIA1 mutants. We use our established seeding method for the wild-type TIA1-LC domain^21^ to obtain homogenous fibrils for each mutant. Transmission electron microscopy (TEM) imaging and Thioflavin T (ThT) assays report on fibril morphology and beta-strand content. Solid state NMR measurements used as a spectroscopic fingerprint allow comparison with wild-type fibrils without complete high-resolution structure determination. The results reveal subtle yet distinct influences of each mutant on the fibril conformation of the TIA1-LC.

## Materials and Methods

Most methods for protein purification and fibril formation follow the same procedure as outlined in detail reference ^21^, and are restated in detail below.

### TIA1-LC A381T, V294M, V360M, P362L, M334I site-directed mutagenesis

This protocol was adapted from our previous work from chapter 3.^21^ Five His-tagged TIA1-LC mutant plasmids were prepared using site directed mutagenesis and the forward and reverse primers with the included mutation. (A381T: 5’-ctgggtttcataccctgtcactcgatacccagaag-3’ and 5’-cttctgggtatcgagtgacagggtatgaaacccag-3; V294M: 5’-caatttgattctgctgttgcatgggatttatgaattccggatc-3’ and 5’-gatccggaattcataaatcccatgcaacagcagaatcaaattg-3’; V360M: 5’-gaggcggttgcattccataatttggtcccatcca-3’ and 5’-tggatgggaccaaattatggaatgcaaccgcctc-3’; P362L: 5’-ccattttgcccttgaggcagttgcactccataatttg-3’ and 5’-caaattatggagtgcaactgcctcaagggcaaaatgg-3’; M334I: 5’-tgcctggccatatattccatatgcaggaacttgc-3’ and 5’-gcaagttcctgcatatggaatatatggccaggca-3’; DNA Technologies). A polymerase chain reaction (PCR) was performed by gently mixing 0.5 µL of the pHis-parallel wild-type TIA1-LC domain plasmid (123.1 ng/µl), 10 µL 5× Phusion GC buffer, 1 µL of 10 mM deoxynucleotides, 1.5 µL of dimethyl sulfoxide, 0.5 µL of Phusion DNA polymerase, 2.5 µL of 10 µM forward primer, 2.5 µL of 10 µM reverse primer into 31.5 µL of sterile water for a total volume of 50 µl. Polymerase, buffer, and dNTPs used were obtained from New England BioLabs. A Bio-Rad T100 thermal cycler was used to perform the PCR with an initial denaturation step at 98 °C for 30 s followed by 30 cycles of 7 s denaturation at 98 °C, 20 s of annealing at 72 °C, and extension for 2 min 42 s at 72 °C, and a final extension for 8 min at 72 °C. PCR reaction vial was taken off the thermal cycler and 1 µL of DpnI (New England Biolabs) was added. The vial was incubated at 37 °C for 1 h and then stored at 4 °C. A Monarch PCR & DNA cleanup kit (5 µg) was used to purify the PCR product (New England BioLabs). The PCR product was then transformed into chemically competent DH5α *E. coli* cells. For the transformation, 50 µL DH5α *E. coli* cells were thawed on ice for 10 min before adding 4 µL of the PCR product. The cell mixture was flicked gently 4-5 times to mix before incubating on ice for 15 min. It was then heat shocked at 42 °C for 90 s and was incubated immediately on ice for 5 min. Afterwards, 600 µL of Luria broth media was added and allowed to grow in incubator shaker at 37 °C, shaking at 220 RPM for 0.5 – 1 h. The cells were streaked onto three agar plates pipetting 25 µL, 75 µL, and 150 µL respectively for each mutant for optimal growth and the plates were left to incubate overnight for ∼17 h at 37 °C. The agar plates were allowed to sit at room temperature before a colony was picked and pipetted into 10 mL of Luria broth with 100 µg/mL ampicillin in a loosely capped 50 mL conical tube. The cells were grown overnight for 17 h, shaking at 220 RPM at 37 °C. A Qiagen QIAprep Spin Miniprep kit was used to purify the amplified plasmid and the sequence for each mutant was confirmed by Sanger sequencing (Genewiz).

### TIA1-LC G355R and E384K mutants

Modification of the TIA1-LC pHis-parallel plasmid for the G355R and E384K TIA1-LC mutant was done by GenScript. The transformation and plasmid amplification were performed using the exact protocol as the other five mutations. All sequences were confirmed by Sanger sequencing (Genewiz).

### Protein Expression and purification

N-terminal His-tagged (MSYYHHHHHHDYDIPTTENLYFQGAMDPEF) ^13^C and ^15^N labeled wild-type and disease mutant TIA1-LC domains (residues I291–Q386) were recombinantly produced using a pHIS-parallel plasmid.^22^ The plasmids were transformed into BL21(DE3) *E. coli* cells and grown in a shaker incubator at 37 °C and 220 revolutions per minute (RPM).

For all unlabeled protein, Luria broth media was used in the presence of 100 µg/ml ampicillin to grow bacterial cultures to an optical density (OD) of 0.6–1.0 at 600 nm measured using a cuvette with a 1 cm pathlength for ∼3 h. Protein expression was induced afterwards by the addition of 0.5 mM isopropyl β-D-thiogalactopyranoside (IPTG) and the cultures were grown in the shaker incubator for 3 - 3.5 h. The cells were centrifuged at 6,000 g for 15 min to harvest. The cell pellets that were harvested were flash frozen using liquid nitrogen before storage at ™80 °C until purification.

The ^13^C, ^15^N labeled proteins were recombinantly expressed in Luria broth media in the presence of 100 µg/ml ampicillin. The cells were grown for ∼3 h to an optical density at 600 nm of 0.6–1.0, measured using a 1 cm pathlength cuvette. The cells were transferred into 1 L of M9 minimal media (45.8 mM Sodium phosphate dibasic heptahydrate, 22.0 mM Potassium phosphate monobasic, 8.6 mM sodium chloride, 2.0 mM magnesium chloride, 0.1 mM calcium chloride, and 100 µg/ml ampicillin 2.0 g of U-^13^C^6^ D-glucose, and 1.0 g of ^15^N ammonium chloride (Cambridge Isotope Labs)) and induced with isopropyl β-D-thiogalactopyranoside (IPTG) to 0.5 mM after incubation of cells for 0.5 h longer. After 3 h the cells were harvested, and flash frozen in liquid nitrogen. All the cells were stored at ™80 °C until purification.

The wild type and disease mutants of TIA1-LC were purified using the same procedure. Cell pellets from protein expression were thawed on ice for ∼15 min. The pellet was resuspended in 40 mL of 6 M guanidinium hydrochloride, 50 mM tris(hydroxymethyl)aminomethane (Tris) pH 7.5, 500 mM sodium chloride, and 1% v/v Triton X-100 along with 4 pellets of Mini EDTA-free Pierce Protease Inhibitor (ThermoFisher Scientific), and 0.25 mg/ml hen egg white lysozyme. Resuspended cells were sonicated using a Branson 250 Sonifier equipped with a 1/4’’ microtip for 20 min in an ice-water bath.

The lysed cells were centrifuged to remove any insoluble material. TIA1-LC was purified using 5 mL of fresh Ni2^+^ resin (Bio-Rad Nuvia IMAC Ni-Charged Resin) equilibrated with 500 mM sodium chloride, 6M urea, and 20 mM 4-(2-hydroxyethyl)-1-piperazineethanesulfonic acid (HEPES) pH 7.5. The clarified lysate was mixed with the resin by rotating in a 50 mL conical tube for 30 min at room temperature. The resin lysate mixture was then added into a glass gravity column with a 2.5 cm diameter (Bio-Rad Econo Column), and the flow through collected. Ten mL of 500 mM sodium chloride, 6M urea, and 20 mM 4-(2-hydroxyethyl)-1-piperazineethanesulfonic acid (HEPES) pH 7.5 was used to equilibrate the column. Then the column was washed with 100 mL of 500 mM sodium chloride, 6 M urea, 20 mM imidazole, and 20 mM HEPES, pH 7.5. Lastly, the column was eluted with 25 mL of 500 mM sodium chloride, 6 M urea, and 20 mM HEPES, pH 7.5. Purified protein was eluted and collected into 1 mL aliquots for storage at ™80 °C. SDS-PAGE with Coomassie staining was used to confirm the purity of the TIA1-LC protein.^21^

### Fibril seed preparation of A381T, V294M, V360M, P362L, M334I, G355R TIA1-LC

The seeding procedure summarized here is taken from the methods outlined in Chapter 2, section 2.3 Materials and Methods under the ‘seeding’ section. Purified TIA1-LC protein was thawed to room temperature and centrifuged for 20 min at 21,300 g to pellet any preformed aggregates. One 300 µL aliquot of 138 µM protein, three 1 mL aliquots of 34.5 µM protein, and one 3 mL aliquot of 34.5 µM protein were immediately prepared using 500 mM sodium chloride, 6 M urea, 200 mM imidazole, and 20 mM HEPES, pH 7.5 to dilute each aliquot to the respective concentrations. Aliquots that were not immediately used once prepared, were frozen and stored at -80 °C to prevent further aggregation. The aliquot of 300 µL at 138 µM protein was concentrated at 14,000 g in 500 µL 3 kDa molecular weight cut off (MWCO) spin concentrator until the volume was less than 100 µL. The purpose of the protein concentration is to eventually have less than ∼0.5 % of the original purification buffer by diluting the concentrated protein multiples times using the spin concentrator. The concentrated protein was then diluted with 300 µL 20 mM HEPES pH 7.5 and 200 mM NaCl, and the protein was concentrated to the same volume for a second time. An additional 300 µL of 20 mM HEPES pH 7.5 and 200 mM NaCl buffer was added into the spin concentrator and the sample harvested into a 1.5 mL tube. The harvested protein was then tip sonicated using a 1/8” tip in cycles of 0.1 s on, 1 s off, 1 min total on time at 10% power (these exact conditions were used for all sonication during the seeding method). Once sonicated, the protein was left on the benchtop for 2 days to quiescently aggregate at room temperature. During the incubation period, the protein (‘seeds 1’) was tip sonicated 1 – 2 times with the exact same conditions.

A protein aliquot of 1 mL at 34.5 µM was thawed out, centrifuged for 20 min at 21,300 g, and dialyzed in 6.4 mm flat-width tubing with a MWCO of 6 to 8 kDA for 4 – 24 h into 1 L of 20 mM HEPES pH 7.5. The protein was harvested from dialysis and spun for 20 min at 21,300 g to remove any preformed aggregates from dialysis. The protein concentration of the supernatant was recorded (∼20.7 µM) by taking an absorbance reading at 280 nm. The ‘seeds 1’ left on the benchtop were tip sonicated with the same previous conditions. An aliquot of 100 µL of the ‘seeds 1’ was transferred into a new 1.5 mL tube, diluted with 300 µL of 20 mM HEPES pH 7.5, and tip sonicated for a second time. This new ‘seeds’ aliquot was then gently pipetted into the centrifuged supernatant from the dialysis as the first round of the seeding process of the TIA1-LC fibrils referred to as ‘seeds 2’. The seeded protein or ‘seeds 2’ was left on the benchtop for at least 24 h to quiescently fibrilize at room temperature. The second protein aliquot of 1 mL at 34.5 µM was thawed out, centrifuged for 20 min at 21,300 g, and dialyzed in 6.4 mm tubing with a MWCO of 6 to 8 kDa for 4 – 24 h into 1 L of 20 mM HEPES pH 7.5. The protein was harvested from dialysis and spun for 20 min at 21,300 g to remove any preformed aggregates from dialysis. The protein concentration of the supernatant was recorded (∼20.7 µM) by taking an absorbance reading at 280 nm. The ‘seeds 2’ left on the benchtop were tip sonicated with the same previous conditions. An aliquot of 100 µL of the ‘seeds 2’ was then transferred into a 1.5 mL tube, diluted with 300 µL of 20 mM HEPES pH 7.5, and tip sonicated. This new ‘seeds’ aliquot was then gently pipetted into the centrifuged supernatant from the dialysis as the second round of the seeding process of the TIA1-LC fibrils referred to as ‘seeds 3’. The seeded protein or ‘seeds 3’ was left on the benchtop for at least 24 h to quiescently fibrilize at room temperature. The exact same steps were followed for the third 1 mL aliquot of protein at 34.5 µM for dialyzing and centrifuging the protein. An aliquot pf 100 µL of the ‘seeds 3’ were diluted with 300 µL of 20 mM HEPES pH 7.5 and tip sonicated. The sonicated and diluted ‘seeds 3’ was added into the dialyzed and centrifuged supernatant. The newly seeded fibrils or ‘seeds 4’ were left on the benchtop for at least 24 h to quiescently fibrilize at room temperature.

Finally, the same steps above to prepare ‘seeds 3’ for thawing, centrifuging, and dialyzing the 3 mL aliquot of protein at a concentration of 34.5 µM was followed. An aliquot of 100 µL of ‘seeds 4’ was diluted with 300 µL of 20 mM HEPES pH 7.5 and tip sonicated. The sonicated aliquot was added into the dialyzed and centrifuged supernatant to fibrilize quiescently on the benchtop at room temperature for at least 24 h. After ∼24 h, the fourth and final round of the seeded fibrils were aliquoted into 1.5 mL tubes in 300 µL fractions and stored at -80 °C until use as seeds. The aliquots stored are used for the seeding step of preparing seeded NMR TIA1-LC fibril samples. An aliquot of each process was set aside for TEM and prepared to monitor seed formation, fibril formation, and homogeneity of each seeded round of fibrils.

### Fibril seed preparation of E384K

Purified protein was thawed to room temperature and centrifuged for 20 min at 21,300 g to pellet any preformed aggregates. To prepare the seeds, an aliquot of 300 µL of protein at 138 µM was diluted with 500 mM sodium chloride, 6 M urea, 200 mM imidazole, and 20 mM HEPES, pH 7.5. The diluted aliquot was concentrated at 14,000 g in a 500 µL spin concentrator with a 3 kDa MWCO until the volume was less than 100 µL. The flow through was discarded and the protein was then diluted with 300 µL of 20 mM HEPES pH 7.5 and 200 mM NaCl. The diluted protein was concentrated in the same spin concentrator to the same volume of less than 100 µL for a second time. The concentrated protein was diluted using 300 µL of 20 mM HEPES pH 7.5 and 200 mM NaCl buffer by adding the buffers into the spin concentrator. The protein was then harvested into a 1.5 mL tube. The harvested protein was then tip sonicated using a 1/8” tip in cycles of 0.1 s on, 1 s off, 1 min total on time at 10% power. Once sonicated, the protein was left on the benchtop for 9 days to quiescently aggregate at room temperature. During the incubation period, the protein or ‘seeds’ was tip sonicated 1-2 times with the exact same conditions.

### TIA1-LC mutant and wild-type NMR seeded fibril samples

For each A381T, V294M, V360M, P362L, and M334I mutant, ∼12 – 15 mg of purified ^13^C, ^15^N labelled TIA1-LC protein was thawed from -80 °C. The protein was diluted to 83 µM using 500 mM sodium chloride, 6 M urea, 200 mM imidazole, and 20 mM HEPES, pH 7.5. The protein was dialyzed for 4 h in 1 L of 20 mM HEPES, pH 7.5 to remove denaturants left from the purification. The protein was then harvested and centrifuged for 20 min at 21,300 g to remove any preformed aggregates. A 300 µL aliquot of the corresponding seeds (see *Fibril seed preparation of A381T, V294M, V360M, P362L, M334I, G355R TIA1-LC* of methods) stored at -80 °C was thawed and tip sonicated using a 1/8” tip in cycles of 0.1 s on, 1 s off, 1 min total on time at 10% power. The sonicated seeds were added into the dialyzed and centrifuged protein. The seeded fibrils were left quiescently on the benchtop at room temperature for a minimum of 7 days. The seeding process was the same for both the wildtype and mutants apart from the G355R and E384K TIA1-LC mutants.

### G355R TIA1-LC NMR fibril sample

∼12 – 15 mg of purified ^13^C, ^15^N labelled G355R TIA1-LC protein was thawed from -80 °C. The protein was diluted to 83 µM using 500 mM sodium chloride, 6 M urea, 200 mM imidazole, and 20 mM HEPES, pH 7.5. The protein was dialyzed for 4 h in 1 L of 20 mM HEPES, pH 7.5 to remove denaturants left from the purification. A 10 µL aliquot of the dialyzed protein was set aside for brightfield measurements of liquid droplets (See *Brightfield microscopy* section of methods). The rest of the protein harvested from dialysis was then harvested and centrifuged for 20 min at 21,300 g to remove any preformed aggregates. The centrifuged pellet was stored at 4 °C for 24 h. The sample was seeded following the protocol above (see *Fibril seed preparation of A381T, V294M, V360M, P362L, M334I, G355R TIA1-LC*). After 24 h, the seeded fibrils were tip sonicated using 1/8” tip in cycles of 0.1 s on, 1 s off, for a total of 30 s on time at 10% power. The centrifuged dialysis pellet stored at 4 °C contained most of the protein and was redissolved using the sonicated seeded fibrils. The redissolved pellet was tip sonicated a second time with the same conditions before leaving it to rotate at room temperature. After 8 days, the fibrils were tip sonicated using a 1/8” tip in cycles of 0.1 s on, 1 s off, for a total of 1 min on time at 10% power and left to rotate at room temperature for 2 days. After 2 days, the rotating fibrils were then spun for ∼2 h at 233,000 g at 12 °C. The supernatant was decanted, and the pelleted fibrils were redissolved in 18 mL of 20 mM HEPES with 0.1 % sodium azide to buffer exchange. The fibrils were tip sonicated using a 1/8” tip in cycles of 0.1 s on, 1 s off, 30 s total on time at 10% power to prepare for ssNMR rotor packing of the sample.

### E384K TIA1-LC NMR fibril sample

∼12 – 15 mg of purified ^13^C, ^15^N labelled E384K TIA1-LC protein was thawed from -80 °C. The protein was diluted to 83 µM using 500 mM sodium chloride, 6 M urea, 200 mM imidazole, and 20 mM HEPES, pH 7.5. The protein was dialyzed for 4 h in 1 L of 20 mM HEPES, pH 7.5 to remove denaturants left from the purification. After the dialysis of ∼12 – 15 mg purified protein was harvested, an aliquot of 10 µL of the protein was set aside for brightfield measurements of liquid droplets (See *Brightfield microscopy* section of methods). The rest of the harvested protein was tip sonicated using a 1/8” tip in cycles of 0.1 s on, 1 s off, with a 1 min total on time at 10% power. The seeds for E384K TIA1-LC (see methods section: *Fibril seed preparation of E384K*) were also tip sonicated using a 1/8” tip in cycles of 0.1 s on, 1 s off, with a 1 min total on time at 10% power. The sonicated seeds were added into the sonicated dialysis harvest and left to rotate at room temperature for 8 days. The seeded fibrils were tip sonicated again using a 1/8” tip in cycles of 0.1 s on, 1 s off, with a 1 min total on time at 10% power and left to rotate at room temperature for 2 days. The fibrils were then spun for ∼2 h at 233,000 g at 12 °C. The supernatant was decanted, and the fibril pellets were redissolved in 18 mL of 20 mM HEPES with 0.1 % sodium azide to buffer exchange. The fibrils were tip sonicated using a 1/8” tip in cycles of 0.1 s on, 1 s off, with a 30 s total on time at 10% power and was left to rotate at room temperature for 4 days.

### Transmission Electron Microscopy

This protocol was taken from our previously published work (from chapter 3, section 3.3 materials and methods).^21^ Five µL of the TIA1-LC domain solutions were incubating for 2 min on Cu or Au 300–400 mesh lacey carbon grids with ultrathin carbon films (Ted Pella). The solution was blotted using a lab tissue, and the grid was washed twice with 5 µL of ultrapure water incubated for ∼10 s, blotting away the water with lab tissue between each wash. The grid was negatively stained using 5 µL of 3% (w/v) uranyl acetate, incubated for ∼10 s, and blotted with a tissue. The grid was also air-dried afterwards for ∼2 min. Images were obtained on a FEI Talos 120C electron microscope operating at 120 kV.

### Fluorescence Assay

A concentrated Thioflavin T (ThT) stock solution (>200 µM, Acros Organics) dissolved in ultrapure water was prepared. The dissolved solution was sterile filtered (0.22 µM PES membrane, Millipore) and the working concentration of 40 µM ThT in 20 mM HEPES pH 7.5 buffer was prepared. The absorbance at 412 nm and an extinction coefficient of 36,000 M^-1^cm^-1^ was used to determine the ThT concentration. The ThT solution was wrapped in foil and stored at 4 °C. All ThT stock solutions were used within 7 d after being prepared.

For each fluorescence measurement of the protein, 50 µL of 40 µM Thioflavin T (ThT) in 20 mM HEPES pH 7.5 buffer and 50 µL of protein was added into a 96-well black polystyrene plate (Fisher). The fluorescence was measured using a BioTek Synergy H1 plate reader with initial linear shaking for 30 s at 1096 cpm (1mm), gain of 100, read height of 5.5 mm, excitation at 440 nm, emission from 460 nm – 620 nm, and with a normal read speed. For the negative control for each measurement, 50 µL of 40 µM Thioflavin T (ThT) in 20 mM HEPES pH 7.5 buffer and 50 µL of 20 mM HEPES pH 7.5 buffer was used. Each measurement was done in triplicate.

Bound ThT assays to monitor fibril formation and the percentage of ThT bound to the fibril were done using the samples from the well plates. The absorbance at 412 nm was measured to determine initial fluorescence values. The protein and control samples were spun for 1 h at 233,000 g at 12 °C. The absorbance at 412 nm of the supernatant was measured. The percentage of ThT bound was measured by subtracting the ThT absorbance at 412 nm of before (the initial reading) and after centrifugation. All measurements were done in triplicate apart from the A381T TIA1-LC mutant, which had only enough protein for one trial.^21^

### Brightfield microscopy

Three µL of the dialysis harvest aliquot set aside from the fibril prep were imaged (see *G355R TIA1-LC NMR fibril sample* and *E384k TIA1-LC NMR fibril sample* methods section). The liquid droplets for both G355R and E384K TIA1-LC mutants were dispensed onto a glass microscope slide. The samples were imaged on Olympus BX51 light microscope with differential interference contrast (DIC) filters, a Diagnostics Instruments RT Slider camera with 6-megapixel sampling, and using a 40× objective lens.^21^

### Solid state NMR data collection

The seeded mutant TIA1-LC fibrils were pelleted for 16-20 h at 233,000 g at 12 °C. The absorbance at 280 nm of the supernatant was measured to calculate any residual soluble protein using a 1 mm pathlength cuvette and the extinction coefficient of TIA1-LC, 46,870 M^-1^cm^-1^ (ProtParam^22^). Using a spatula the centrifuged fibril pellets were transferred into 3.2 mm thin-walled pencil-style zirconia rotors (Revolution NMR). The samples were compacted by spinning the rotors with the samples at 25,000 g for ∼3 h at 8 °C. The rotor drive tip and cap were sealed to the rotor using cyanoacrylate gel. The hydrated samples are approximately 30% protein based off the volume of the NMR rotor (36 µL) and the average protein density (1.35 g/cm^3^).

The ssNMR collection was acquired following the protocol from our previously published work, restated below.^21^ All ssNMR experiments were collected at the UC Davis NMR Campus Core Facility (Davis, CA) and the National High Magnetic Field Laboratory (Tallahassee, FL) on 18.8 T magnets. The probes used for all experiments were the BlackFox NMR and Low-E triple resonance 3.2 mm MAS probes. The sample temperature for all experiments was ∼10 °C. ^1^H decoupling of 83.3 kHz was used for cross polarization experiments. For the ^15^N-^13^C cross polarization steps, CW was used and for the indirect acquisition periods, SPINAL-64 was used. Ten kHz WALTZ-16 ^1^H decoupling was used for INEPT experiments. A dehydrated sample of 1-^13^C labeled Ala powder was used to externally reference the observed chemical shifts to the DSS scale with the Adamantane ^13^C downfield peak at 40.48 ppm. Using a ^1^H-^13^C 1D experiment, a variable length spin-echo period directly after the cross polarization step was inserted for ^13^C T_2_ measurements. Using a 1D NCA experiment, a variable length spin-echo period was inserted between the ^1^H-^15^N and ^15^N-^13^CA cross polarization steps for ^15^N T_2_ measurements. 2D experiments were used for the site-resolved ^15^N T_2_ experiments. High power 83.3 kHz ^1^H decoupling was used for echo periods which ranged up to 10.2 ms for ^13^C and 15.2 ms for ^15^N. TopSpin 3.6 software was used to fit the integrated 1D signal intensities to single exponentials and NMRPipe was used to extract the T_2_ values from the 2D spectra and fit to single exponentials using Python scripts. The contours of all carbon-carbon correlation spectra were plotted with an increasing factor of 1.25 and the nitrogen-carbon correlation spectra with an increasing factor of 1.20. Table 2 includes detailed parameters for each experiment.

## Results

### Seeded fibril formation of TIA1-LC disease mutants

Seven mutations, A381T, V294M, V360M, P362L, M334I, G355R, and E384K, linked to clinical cases of ALS^12^ reside in the TIA1-LC. Figure 1 shows these mutations span the entire TIA1-LC, with one mutant (G355R) in the region known to form the rigid core of TIA1-LC wild type fibrils formed by a repetitive seeding process.^21^ The influence of these sequence changes on the ability of the TIA1-LC to form fibrils was investigated through a repetitive seeding process we previously used to investigate the wild type TIA1-LC.^21^ Figure 2 shows that five of the seven mutants (A381T, V294M, V360M, P362L, and M334I) behaved similarly through the seeding process, forming short ∼50 nm seed fibrils after incubation with moderate salt and repetitive sonication at near neutral pH. Adding these seeds to soluble preparations of the TIA1-LC domain without salt at near neutral pH led to the visually homogenous preparations shown in Figure 2. Table 1 shows the residual soluble TIA1-LC concentrations for each mutant obtained from a sedimentation assay.

**Table 1:** Average soluble protein residual concentrations.

| **TIA1-LC Mutant** | **Average Residual Soluble Protein in No Salt Concentration** |
| --- | --- |
| A381T | 0.9 μM |
| V294M | 0.1 μM |
| V360M | 0.5 μM |
| P362L | 1.4 μM |
| M334I | 0.0 μM* |
| G355R | 0.0 μM* |
| E384K | 0.5 μM |

\*Concentrations below 0.0 μM are not detectable using absorbance at 280 nm with the equipment available.

**Table 2:**
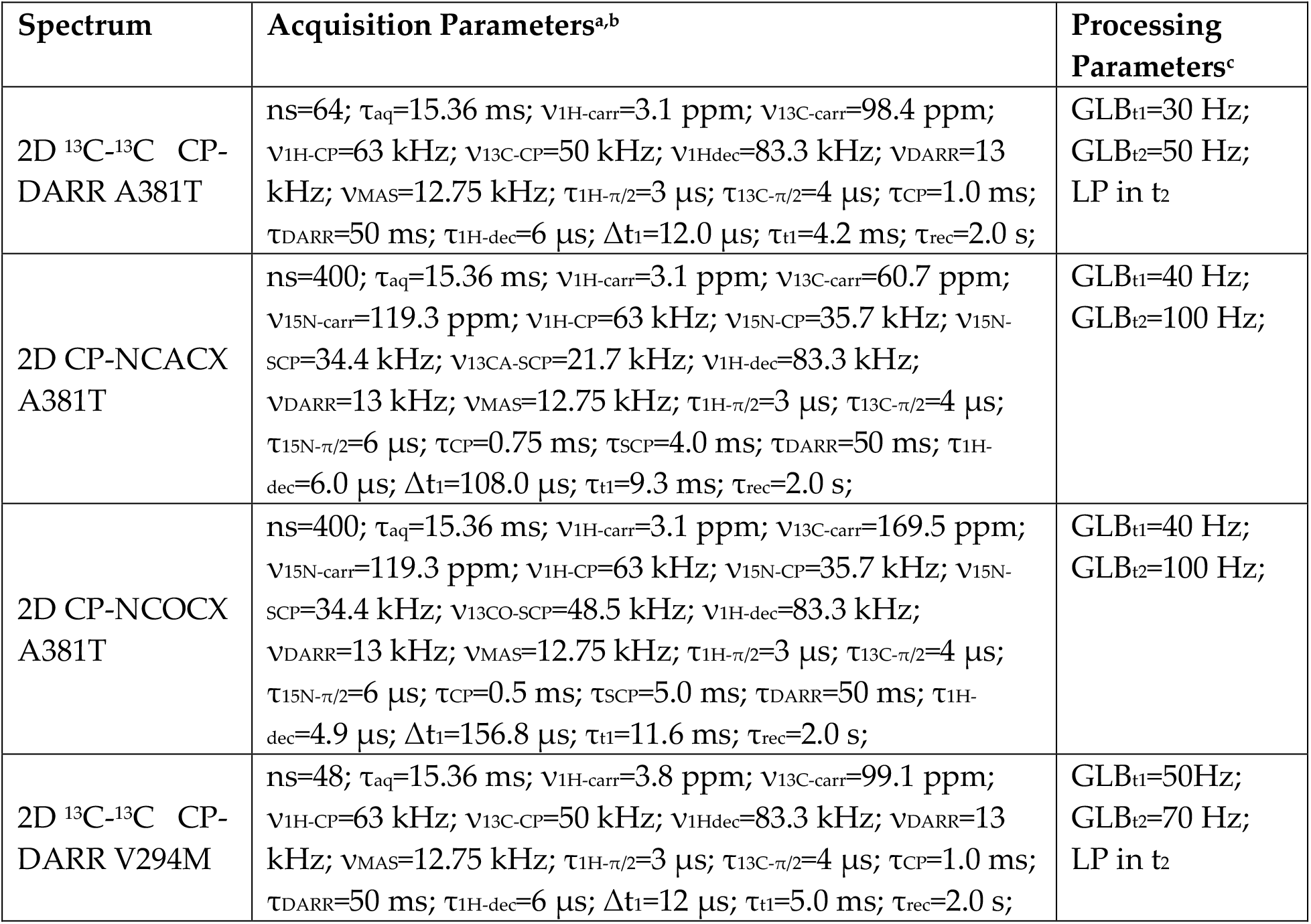
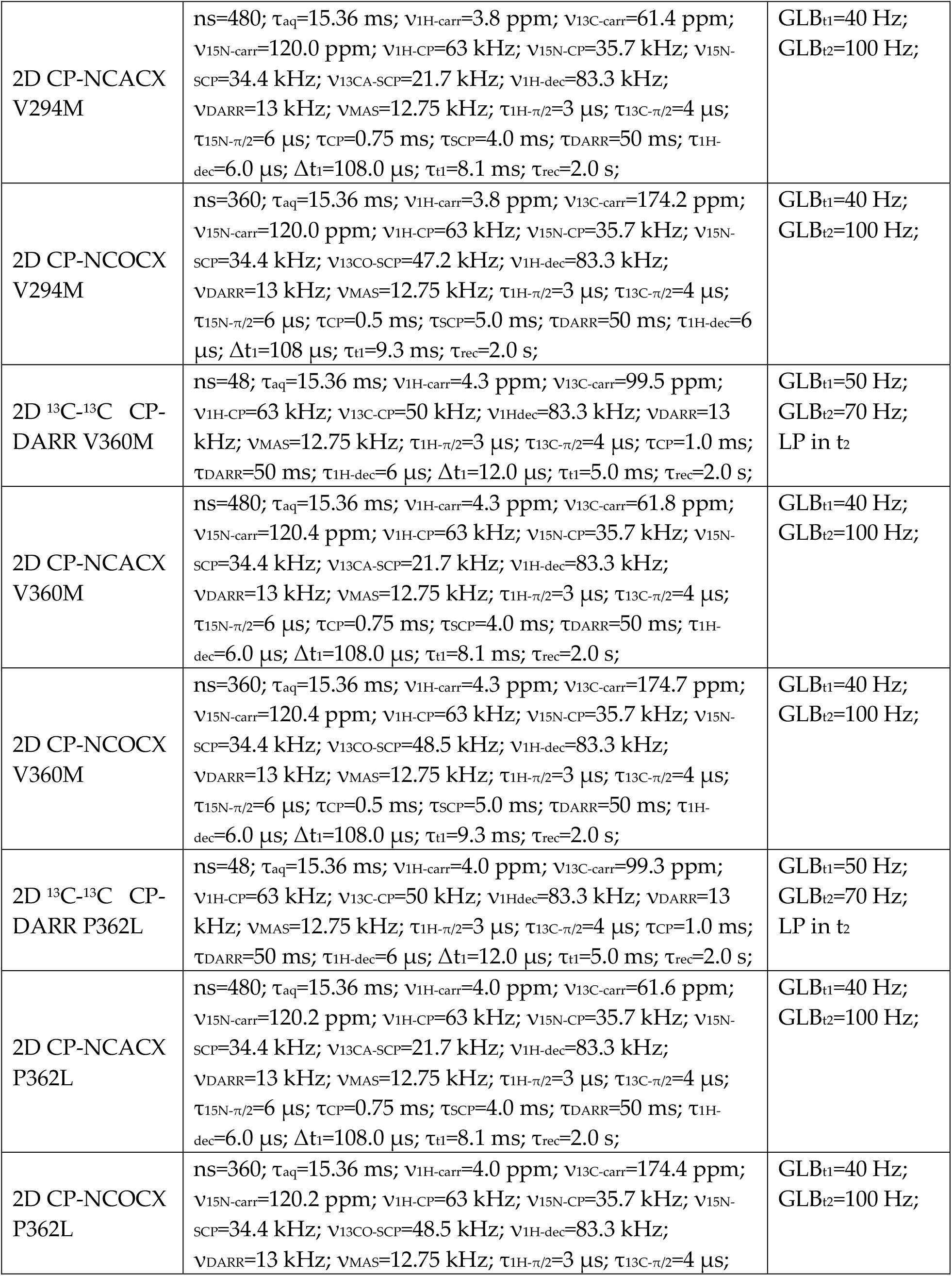
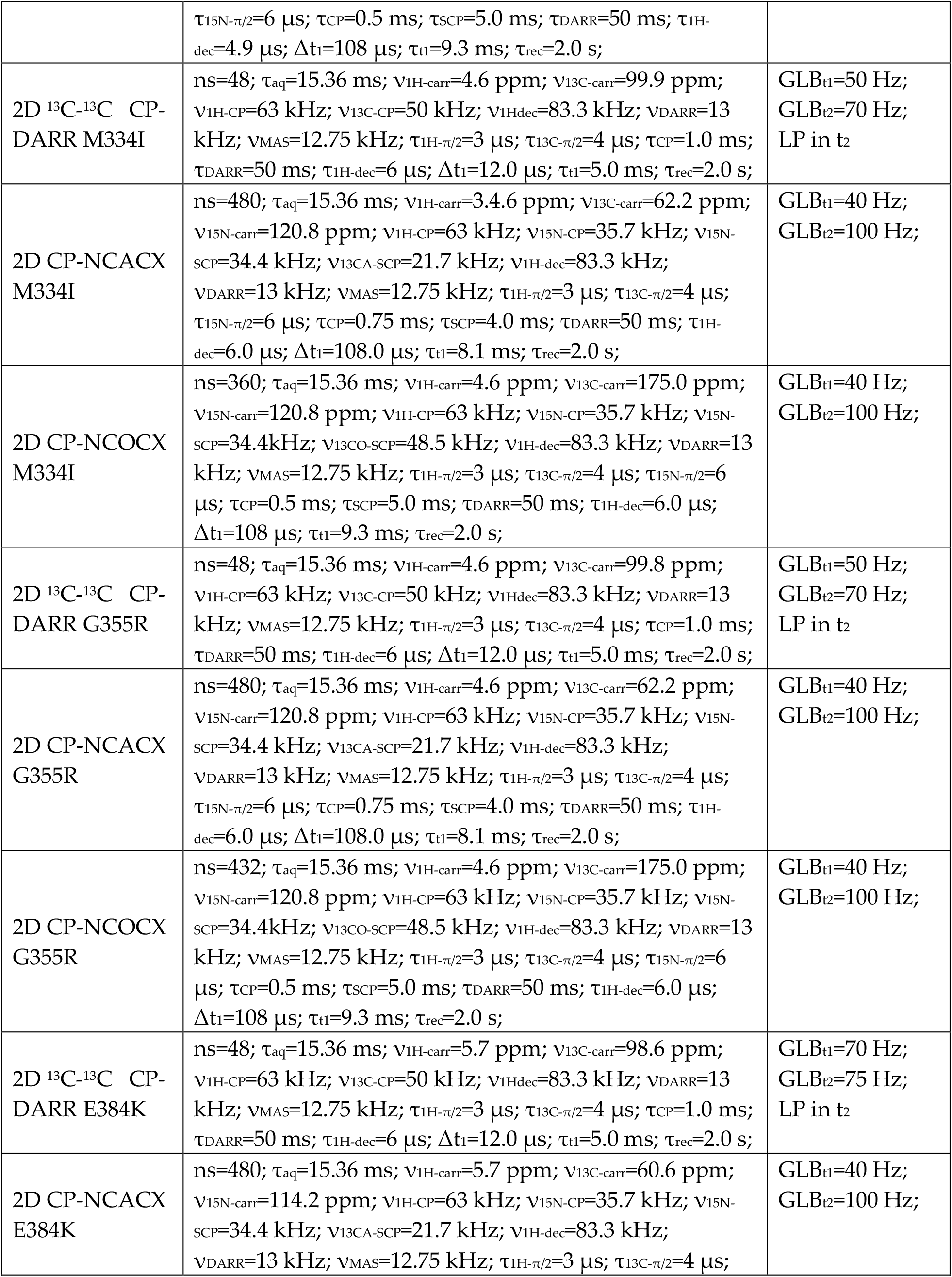
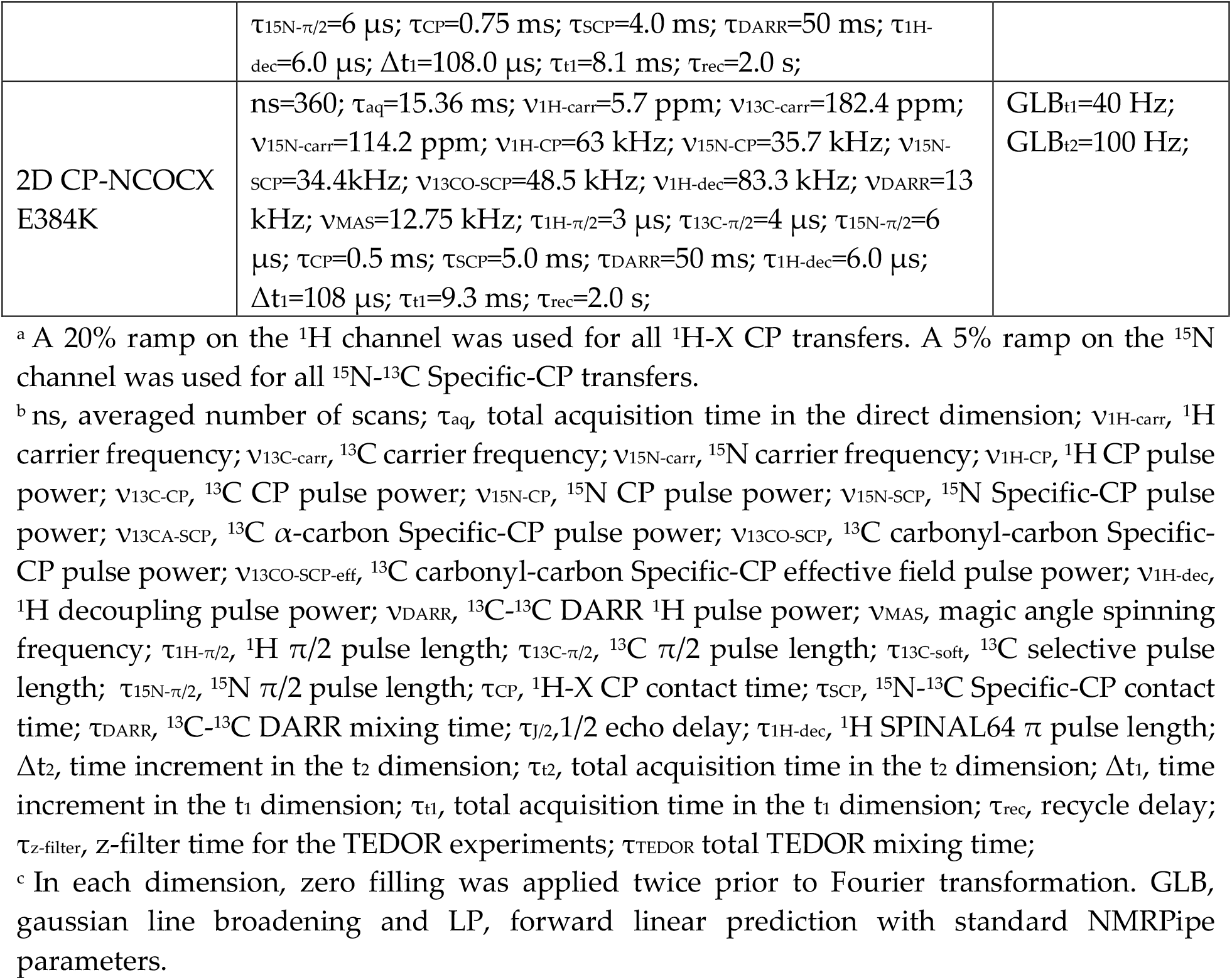
Solid state NMR data collection and processing parameters.

**Figure 1.**
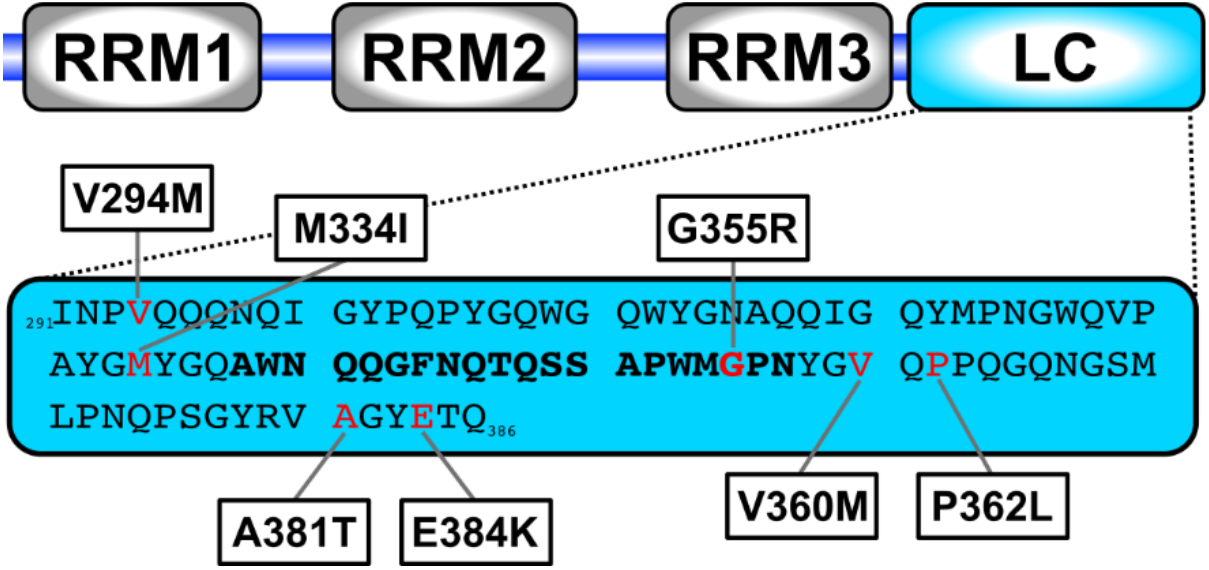
TIA1 domain structure and low complexity domain sequence. RRM are RNA recognition motifs and LC is the low complexity domain. Amino acids in red indicate the location of disease mutations with the specific mutations indicated in the boxes. Bold text highlights the twenty residues comprising the rigid core of the fibrils formed by the wild type LC domain.

**Figure 2.**
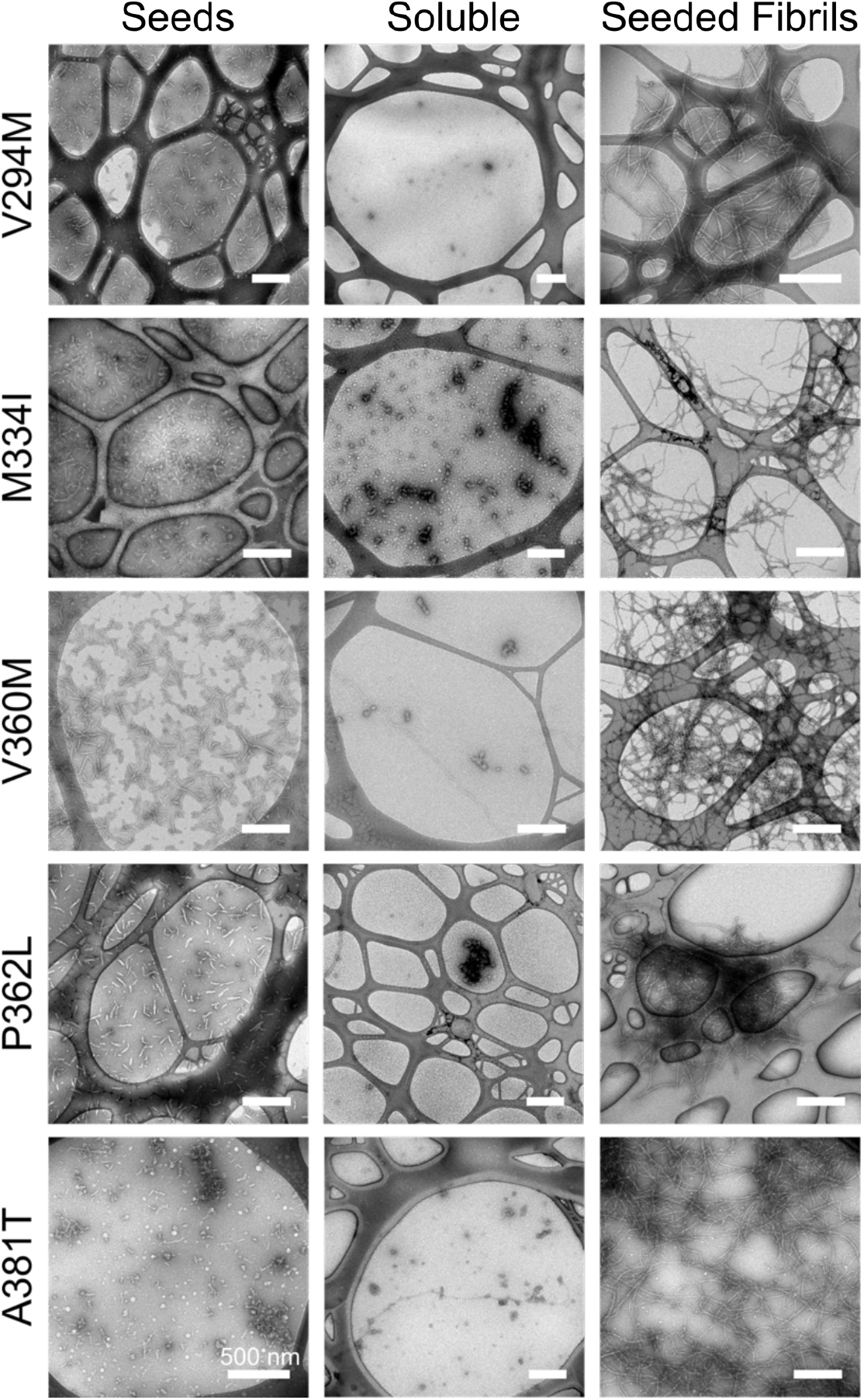
Negative stain TEM images of seeded fibril formation. Left column, mutant TIA1-LC protein seeds. Center column, mutant TIA1-LC protein after dialysis into buffer at pH 7.5 without salt. Right column, mutant TIA1-LC protein after incubation of the soluble protein with no salt with the seeds. Each row represents a different mutation, indicated at the left side of the row.

The G355R and E384K mutant TIA1-LC proteins behaved differently under the same conditions as the other mutants. Immediately upon removal of the denaturant used during purification, the concentrated samples for these mutants in the no salt condition contained the phase separated liquid droplets shown in Figure 3. The other mutants did not form visible phase separations under these conditions. Centrifugation of the G355R mutant to remove phase separated and aggregated protein provided a solution with a concentration high enough (20.7 µM) for seeding. The E384K mutant was much less soluble (6.9 µM) under these conditions and was not amenable to our seeding procedure. The phase separation of G355R and E384K shown in Figure 3 are qualitatively different visually, with the G355R liquid droplets appearing smaller than the droplets formed from the E384K mutant. To keep the sample preparations as consistent as possible, the G355R and E384K proteins were allowed to aggregate in the high salt condition, mechanically processed to form seeds, mixed with TIA1-LC protein prepared without salt but not centrifuged after dialysis, and allowed to incubate for the same time as the other mutants. TEM images of the G355R and E384K preparations are shown in Figure 4.

**Figure 3.**
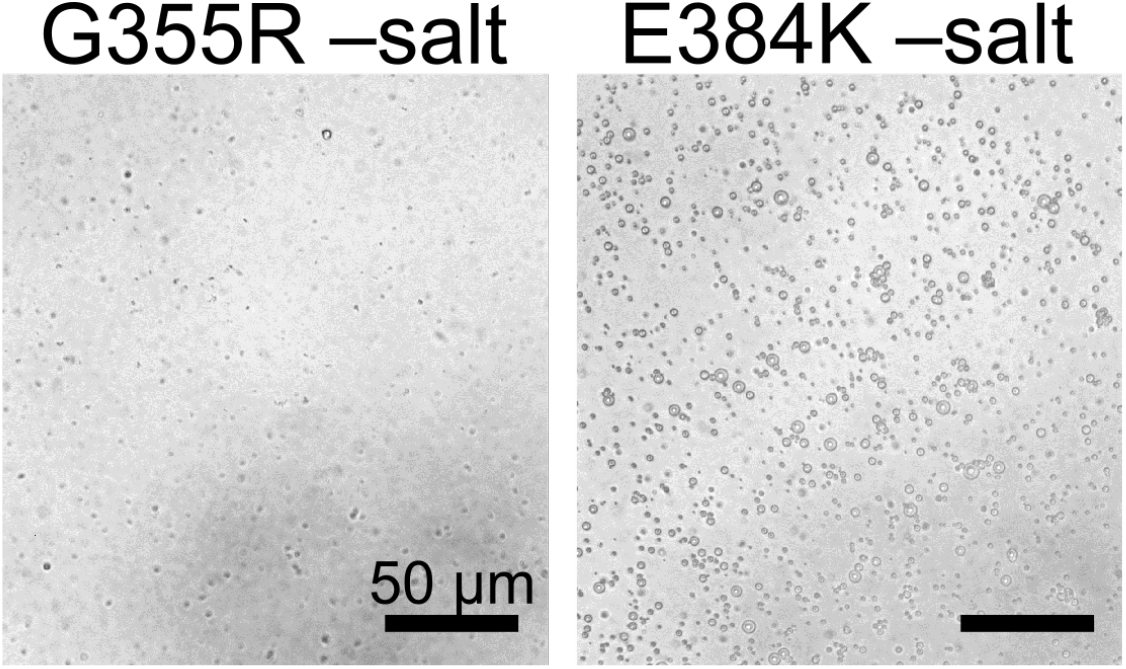
Bright field images of G355K and E384K TIA1-LC. Protein was imaged after dialysis into non-denaturing buffer at pH 7.5 with no salt.

**Figure 4.**
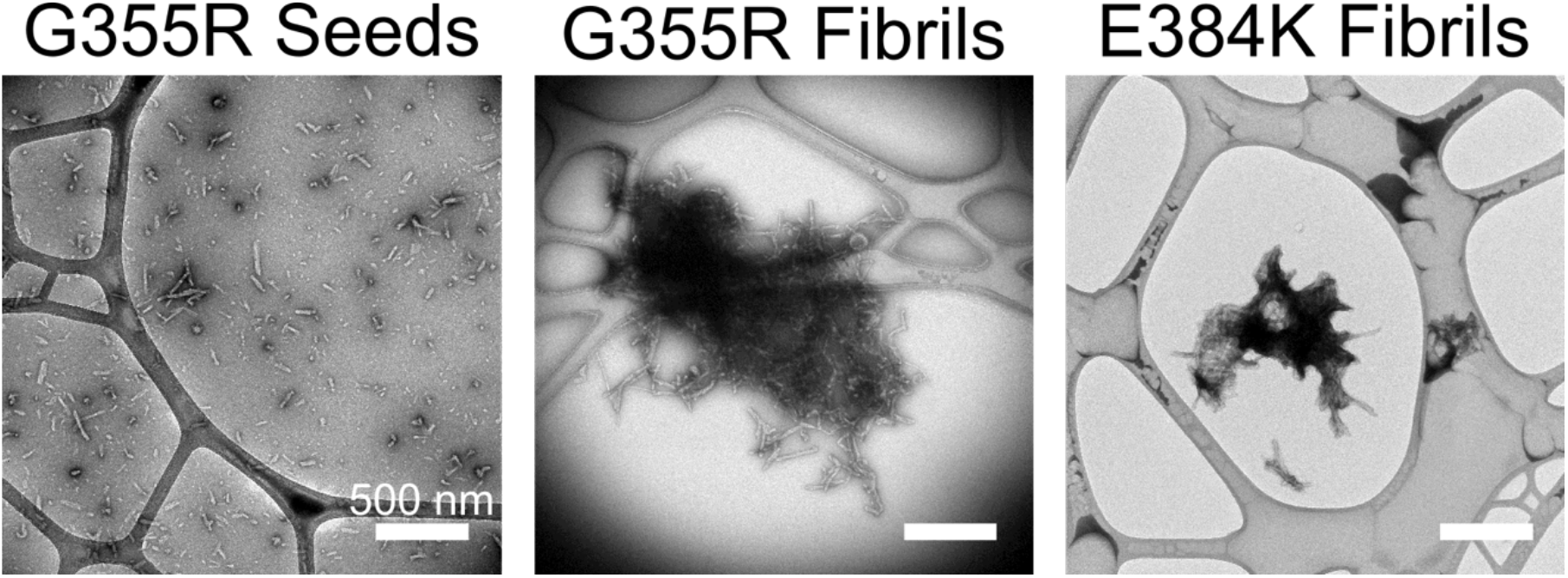
Negative stain TEM images of G355R and E384K TIA1-LC mutant proteins. Left image is the G355R seeds obtained after incubation with 200 mM sodium chloride at pH 7.5 and mechanical processing. Center image is the G355R fibrils obtained after the seeding process. Right image is the E384K protein with no salt at pH 7.5 after incubation with protein aggregated with 200 mM sodium chloride at pH 7.5 and mechanically processed.

### TIA1-LC mutants form β-sheet rich fibrils

Our previous measurements showed a significant increase in thioflavin-T (ThT) fluorescence after incubation with wild type seeded TIA1-LC fibrils.^21^ Figure 4.5A shows that mixing ThT with the seeded fibril preparations for the mutant TIA1-LC constructs revealed similar increases in fluorescence. The increase in fluorescence is consistent with a fibril structure with significant cross-β arrangement of β-strands in the rigid core region.^23^

**Figure 5.**
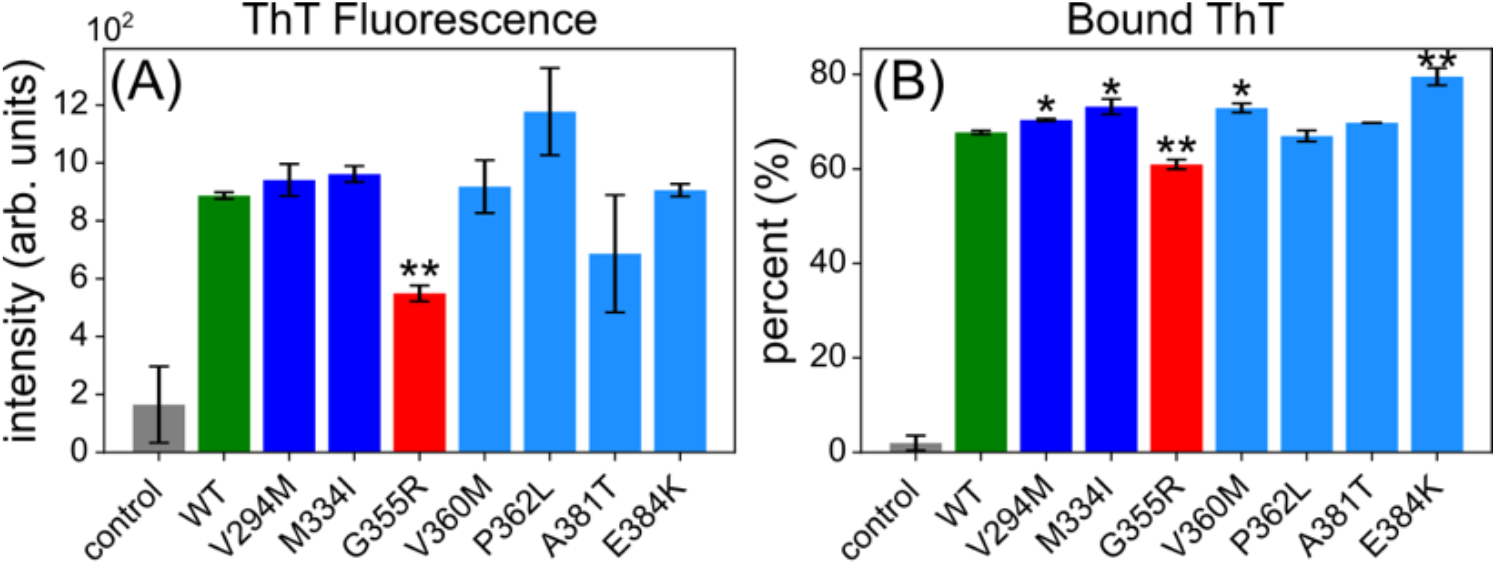
ThT fluorescence measurements on TIA1-LC mutants. (A) ThT fluorescence per bound ThT molecule, calculated by dividing the measured fluorescence signal by the amount of ThT bound to the fibrils. (B) Percent of the total ThT bound to the fibrils. Error bars are +/-the standard deviation. *p<0.05, **p<0.01 by Student’s t-test relative to the wild type TIA1-LC.

Figure 4.5B shows the wild type and mutant TIA1-LC fibril samples all bind >60% of the ThT dye after incubation. The fluorescence data in Figure 4.5A is the intensity per amount of bound ThT, to better report on the similarities and differences in the ThT fluorescence properties of each TIA1-LC construct. Together the data in Figure 4.5A–B show that the wild-type and the five mutants that behaved similarly during the seeding procedure cause similar increases in ThT fluorescence and bind similar amounts of the ThT. These data are therefore consistent with similar β-strand content for the wild type, V294M, M334I, V360M, P362L, and A381T TIA1-LC proteins (p-value less than 0.05 relative to wild type via Student’s t-test). The G355R mutant both binds significantly less ThT and has lower ThT fluorescence intensity (p-value less than 0.01 relative to wild type via Student’s t-test), suggesting a reduced β-strand content for this mutant, or at least reduced accessibility of the ThT molecule to the rigid β-strand core of the fibril. However, while the E384K mutant binds more ThT, it does not cause significantly more fluorescence, consistent with a similar β-strand content as the wild type TIA1-LC.

### Structure of disease mutant TIA1-LC fibrils differ from the wildtype

As shown in Figures 6 and 7, solid state NMR cross polarization-based carbon-carbon and nitrogen-carbon dipolar correlation experiments performed on wild-type and mutant seeded fibril preparations reveal highly similar resonance patterns. Although not obvious at first glance, our previous study on wild-type TIA1-LC fibrils^21^ revealed these spectra likely contain both sharp signals arising from a rigid, homogenous fibril core region and broad signals arising from loosely ordered regions in heterogenous conformations. For spectra like these, where the protein sequence is largely the same (e.g. a single amino acid change), difference spectroscopy has proven useful in distinguishing differences between visually similar spectra. Figure 8 shows the 2D CC and 2D NCACX difference spectra for the each mutant compared to the wild-type. All mutants show a mixture of positive (mutant, variable color) peaks and negative peaks (wild-type, green), indicating that certain regions of the wild-type fibril structure is destabilized by the mutations. The pattern of peaks also vary among all mutants, suggesting each mutant destabilizes the wild-type fibrils differently and that each mutant stabilizes a slightly different conformation for the TIA1-LC. Note these differences are not explained by each sample having a different amino acid substitution. The mutants A381T, V294M, V360M have fewer peaks in the difference spectra than the P362L, M334I, G355R, E384K mutants. The mutants A381T, V295M, and V360M have a lot of shared signals with similar chemical shift values to each other in the difference spectra such as the Gly, Thr, Asn/Asp signals. The M334I and E384K mutants also have similar signals not present in the wildtype spectra.

**Figure 6.**
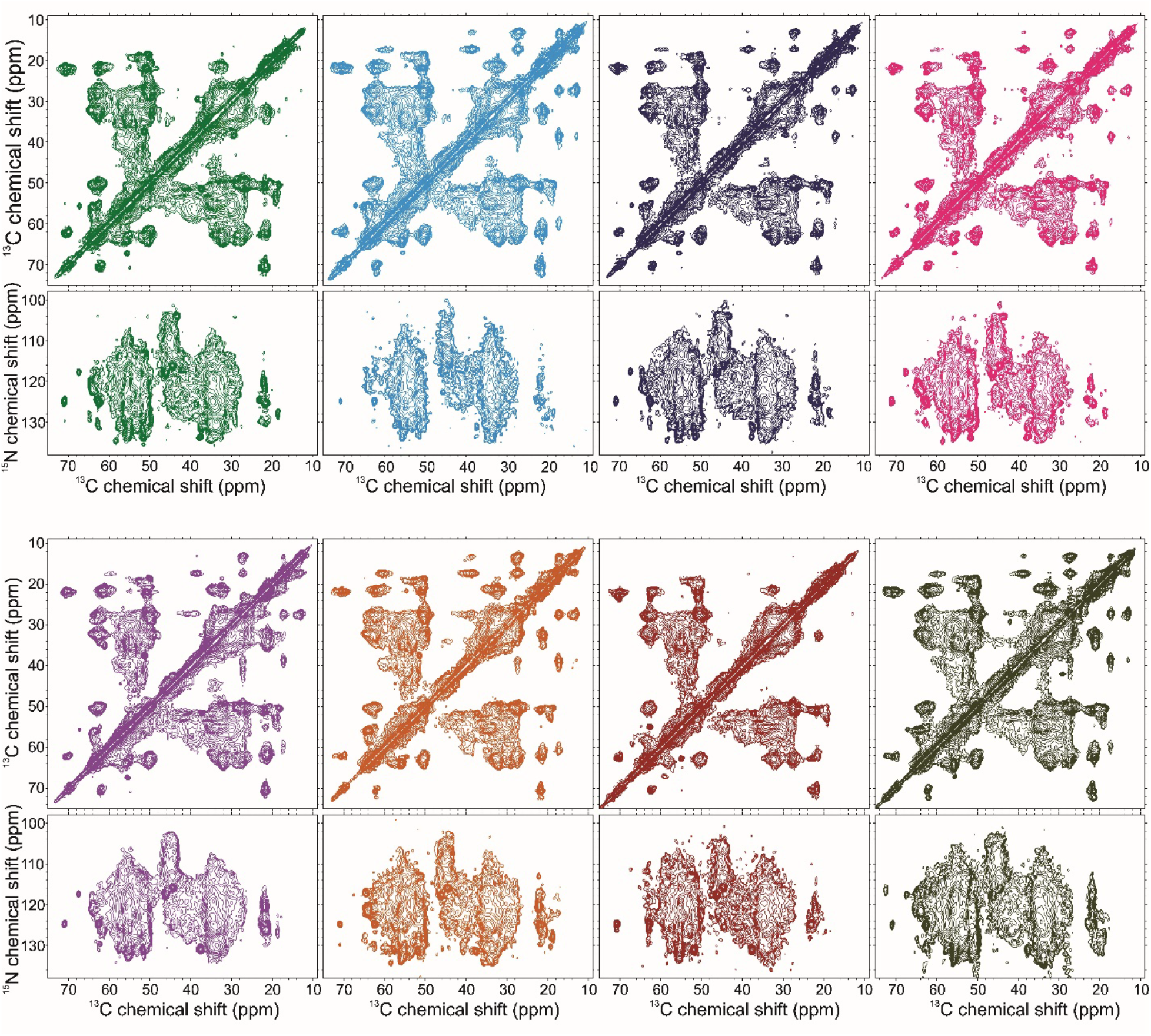
Dipolar-based carbon-carbon and NCACX spectra of WT TIA1-LC and TIA1-LC mutant fibrils. Top from left to right: WT TIA1-LC, A381T TIA1-LC, V294M TIA1-LC, V360M TIA-LC; Bottom from left to right: P362L TIA1-LC, M334I TIA1-LC, G355R TIA1-LC, E384K TIA1-LC.

**Figure 7.**
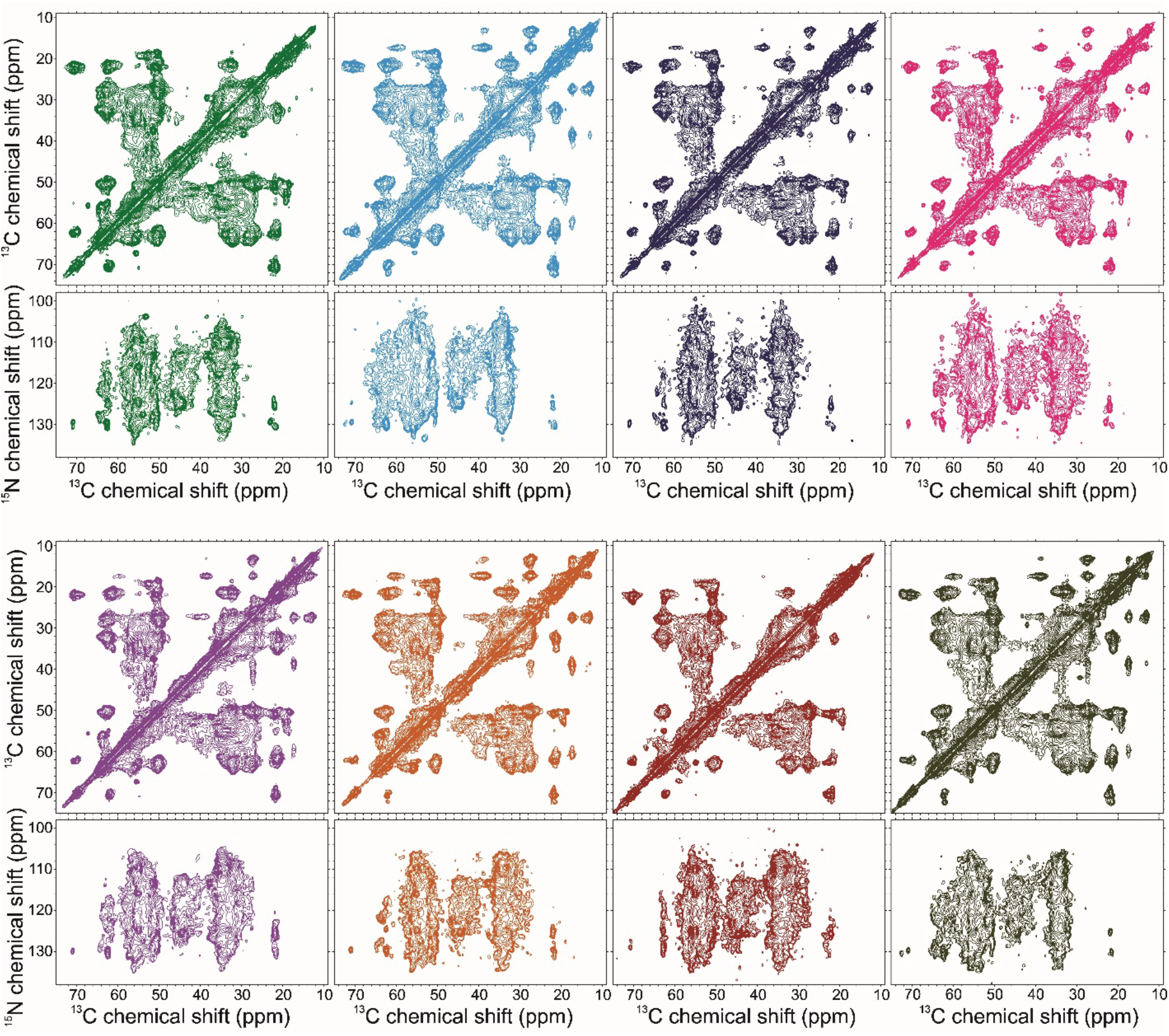
Dipolar-based carbon-carbon dipolar assisted rotational resonance and NCOCX spectra of WT TIA1-LC and TIA1-LC mutant fibrils. Top from left to right: WT TIA1-LC, A381T TIA1-LC, V294M TIA1-LC, V360M TIA-LC; Bottom from left to right: P362L TIA1-LC, M334I TIA1-LC, G355R TIA1-LC, E384K TIA1-LC.

**Figure 8:**
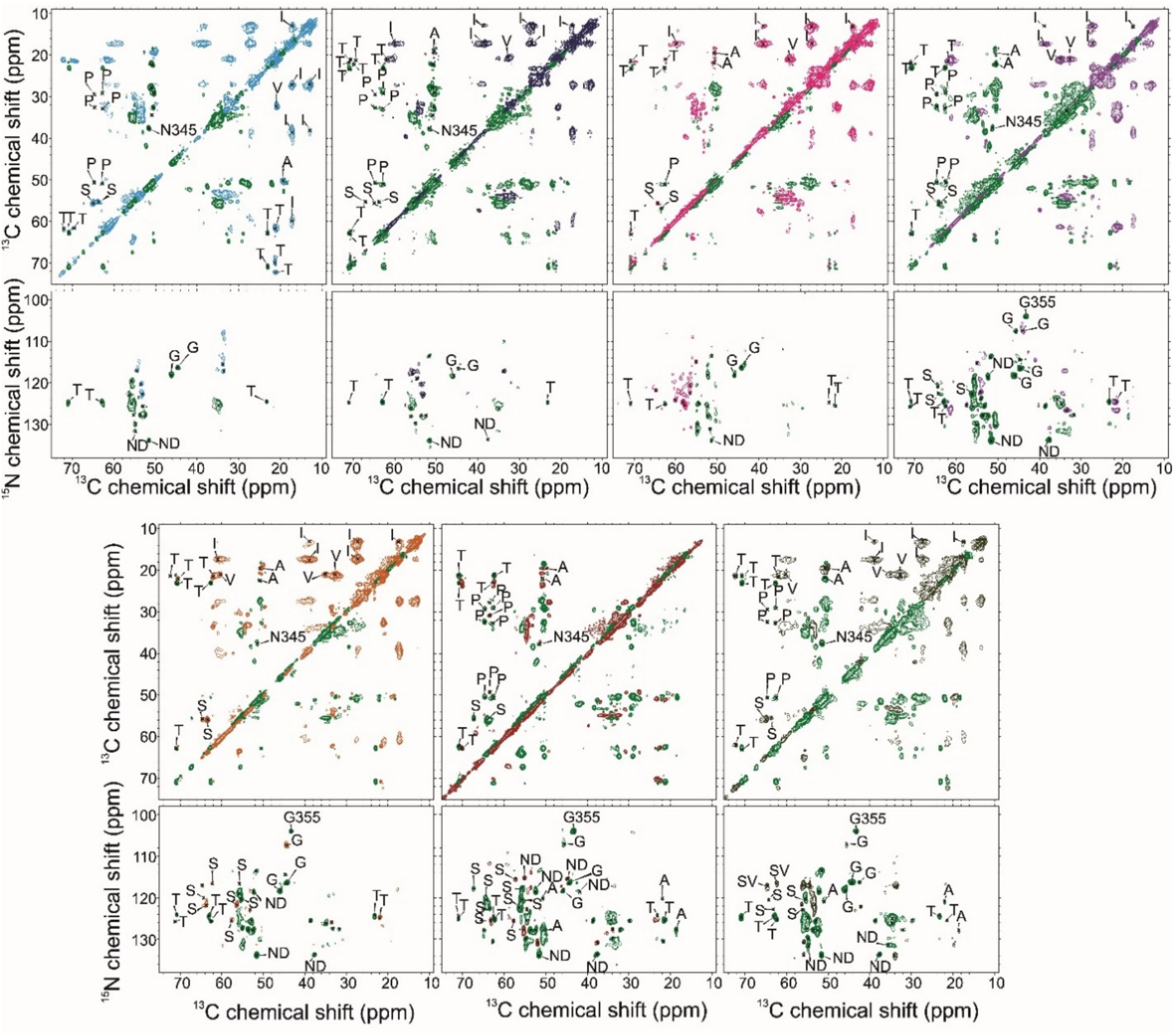
Difference spectra comparing each TIA1-LC mutant to the wild-type TIA1-LC. Top from left to right: A381T TIA1-LC, V294M TIA1-LC, V360M TIA-LC, P362L TIA1-LC; Bottom from left to right: M334I TIA1-LC, G355R TIA1-LC, E384K TIA1-LC. Peaks in green are more present in the wild-type spectrum, and each mutant is represented by a different color. Signals are identified by type where possible and the N345 and G355 signals are unambiguous.

## Discussion

The TIA1-LC disease mutants form fibrils which are structurally different from the wildtype TIA1-LC based on our solid state NMR analysis. Although the spectra all appear visually similar, careful application of difference spectroscopy highlights differences for each of the mutants (Figure 8). Interestingly, none of the mutants completely destabilize the entire structure of the wild-type fibril. The rigid core of the wild-type protein contains 20 amino acids.^21^ Here, none of the mutant difference spectra contain the expected signature for complete destabilization: 20 strong sharp signals in green at precisely the chemical shift values measured for the wild-type protein. Also, there is no obvious new conformation dictated by the mutations, which would manifest as a significant number of strong sharp peaks corresponding to the mutant spectra. All mutant fibrils have similar stabilities in our sedimentation assay (Table 1), with the weakest dissociation constant being 1.6 · 10^−6^, although a statistical comparison is not possible with our current data set. The beta-strand content is only significantly different for the G355R mutant (Figure 5), which is the only mutant in the rigid core of the wild-type fibrils.

Together our data suggest that the ALS-linked mutations in the TIA1-LC domain do not significantly alter the end state of TIA1-LC aggregation. While a secondary fibril forming core could still exist in the TIA1-LC, it does not seem to be triggered by one of these seven ALS mutations. Finally, our results reinforce the primary effect of the mutations being disruption of the condensed state of the protein (i.e. stress granules and liquid droplets) rather than stabilization of the fibrils resulting from rigidification of the condensed state.

## Acknowledgements

This work was fully supported by award R35GM142892 to D.T.M. from the National Institute of General Medical Sciences of the National Institutes of Health. The content is solely the responsibility of the authors and does not necessarily represent the official views of the National Institutes of Health.

